# Two long non-coding RNAs, *SVALKA* and *SVALNA*, regulate *CBF1* and *CBF3* via multiple mechanisms

**DOI:** 10.1101/2025.05.09.653013

**Authors:** Isabell Rosenkranz, Sarah Mermet, Vasiliki Zacharaki, Peter Kindgren

**Affiliations:** Umeå Plant Science Centre, Department of Forest Genetics and Plant Physiology, Swedish University of Agricultural Sciences, 90187 Umeå, Sweden

**Keywords:** Arabidopsis, cold acclimation, epigenetic regulation, long non-coding RNAs

## Abstract

Long non-coding RNAs (lncRNAs) are emerging as key regulatory players of coding gene expression in eukaryotes. Here, we investigate the roles of the lncRNAs *SVALKA (SVK)* and *SVALNA (SVN)* in regulating *CBF1* and *CBF3* gene expression in Arabidopsis under cold stress conditions. We used Native Elongation Transcript Sequencing, CRISPR-Cas9 deletion strategies, and RT-qPCR to analyze the transcriptional dynamics and regulatory mechanisms of *SVK* and *SVN*. Our results demonstrate that *SVK* functions as a *cis*- and *trans*-acting lncRNA, regulating both *CBF1* and *CBF3* through RNAPII collision and chromatin remodeling, while *SVN* serves a *cis* role by negatively regulating *CBF3* via a RNAPII collision mechanism. We identified isoforms of *SVK*, originating from distinct transcription start sites and undergo alternative splicing to adapt structural stability, crucial for their regulatory functions. This study elucidates the complex interplay of lncRNAs in gene regulation, highlighting their essential roles in modulating responses to environmental stresses. Our findings contribute to a deeper understanding of the mechanisms underlying lncRNA functionality and their significance in gene regulatory networks in eukaryotes.

## INTRODUCTION

Long non-coding RNAs (lncRNAs) are a diverse class of RNA molecules longer than 200 nucleotides that play essential roles in gene regulation (Rinn & Chang, 2012; Zhao *et al*, 2018). In contrast to small RNAs, which primarily engage in RNA interference pathways, lncRNAs function through various mechanisms such as acting as molecular scaffolds, decoys, guides, or enhancers involved in transcriptional and post-transcriptional processes (Geisler & Coller, 2013; Kashi *et al*, 2016). They influence key biological functions by interacting with DNA, RNA, and proteins, thereby modulating chromatin dynamics, transcription factor activity, and RNA stability (Csorba *et al*, 2014; Wang *et al*, 2011; Wilusz *et al*, 2009).

In plants, lncRNAs contribute to key developmental processes such as flowering (Csorba *et al*., 2014; Kim *et al*, 2017), hormone signaling (Ariel *et al*, 2014) and responses to abiotic stresses (Gómez-Martínez *et al*, 2023; Kindgren *et al*, 2018). By interacting with chromatin-modifying complexes, such as Polycomb Repressive Complex 2 (PRC2), lncRNAs can direct histone modifications to specific loci, thereby fine-tuning gene expression (Brockdorff, 2013). For instance, at the *FLOWERING LOCUS C* (*FLC*) gene locus in *Arabidopsis thaliana*, several lncRNAs have been proposed to recruit PRC2, via binding of the PRC2 subunit CLF, to deposit the repressive histone mark H3K27me3 (Heo & Sung, 2011; Kim *et al*., 2017). Another lncRNA, *COOLAIR*, functions in parallel, and in a PRC2-independent mechanism, to remove the active H3K36me3 mark from the *FLC* gene body (Nielsen *et al*, 2024). The activity of two distinct repressive mechanisms is believed to give the plant flexibility and temporal plasticity to regulate *FLC* output in response to cold (Nielsen *et al*., 2024).

As sessile organisms, plants must rely on intricate regulatory networks to perceive and respond to fluctuating environmental conditions (Urano *et al*, 2010). A well-characterized pathway involved in cold stress adaptation is the CBF-dependent pathway, in which C-repeat/dehydration-responsive element binding factors (*CBF*s) serve as central regulators for cold acclimation (Gilmour *et al*, 1998; Jaglo-Ottosen *et al*, 1998). In Arabidopsis, the *CBF1-3* genes are arranged in tandem on chromosome 4 and are tightly controlled during cold stress to enhance plant survival (Gilmour *et al*., 1998; Medina *et al*, 1999). Like *FLC, CBF1* and *CBF3* are negatively regulated by lncRNAs. *SVALKA* (*SVK*), an antisense lncRNA transcribed downstream of *CBF1*, fine-tunes the expression of both *CBF1* and *CBF3*, albeit with distinct mechanisms (Gómez-Martínez *et al*., 2023; Kindgren *et al*., 2018). Upon 2 to 4 hours of low-temperature exposure, RNA Polymerase II (RNAPII) read-through transcription of a short *SVK* α isoform leads to head-to-head RNAPII collisions within the *CBF1* gene body. These collisions trigger premature termination of *CBF1* transcription and degradation of the transcript, ultimately repressing *CBF1* expression (Kindgren *et al*., 2018). Under normal growing conditions (22°C), a longer isoform of *SVK* α (*SVK-L*) functions as a natural antisense transcript to *CBF1*, reducing *CBF1* mRNA levels by forming double-stranded RNA that is subsequently degraded (Zacharaki *et al*, 2023).

Additionally, a longer isoform (*SVK* β) originating from a distal TSS of *SVK* interacts with CLF to deposit H3K27me3 marks over the *CBF3* gene body after prolonged exposure to low temperatures (Gómez-Martínez *et al*., 2023). The dynamic RNA regulation of *SVK* and its various isoforms are essential for its regulatory functions, allowing the lncRNA to engage with multiple molecular partners and participate in complex gene networks.

Here, we identify and characterize a novel long non-coding RNA *SVALNA*, transcribed downstream of *CBF3*. Unlike *SVK, SVN* is a target for the nuclear exosome and is rapidly degraded, confining it to a negatively regulated *cis* role on *CBF3* by RNAPII collision in a temporal and mechanistic manner akin to the *SVK-CBF1* circuit. This suggests a complex dynamic where *SVN* serves as a critical regulatory element that fine-tunes *CBF3* expression prior to epigenetic repression by *SVK*. Furthermore, we show that the *trans* role of *SVK* on *CBF3* depends on an epigenetically controlled switch from a proximal to a distal TSS and splicing. Thus, our research highlights the active involvement and temporal regulation of distinct isoforms of lncRNAs to achieve regulatory flexibility in response to unfavorable conditions.

## RESULTS

### Confirmation of the collision mechanism along the *CBF1* gene body

To visualize the collision zone of *CBF1* and *SVK* that occurs post-peak of *CBF1* expression (after three to eight hours at 4°C), we used plant Native Elongation Transcript Sequencing (plaNET-seq) data (Kindgren *et al*, 2020). Accumulation of actively transcribing RNAPII complexes can be seen on both strands designating a hallmark for RNAPII collision (Fig. 1A). To further support the collision mechanism, we used a line that harbors a transfer DNA (T-DNA) insertion in the *CBF1* promoter which greatly limits *CBF1* expression (*cbf1-3*, Fig. 1B) (Zacharaki *et al*., 2023). In T-DNA lines, large inserts of DNA disrupt the genomic context of the insertion site (Alonso *et al*, 2003). We also included the *uns-1* mutant that uncouples the expression of *CBF1* and *SVK*, and the *svk-1* mutant that knocks out *SVK* (Kindgren *et al*., 2018). Both mutations lead to increased *CBF1* expression after cold exposure (Kindgren *et al*., 2018). Our hypothesis was that restriction of RNAPII transcription on one strand should increase the steady state RNA levels of the transcript derived from the other strand due to fewer collision events occurring after cold exposure (Fig. 1B). As expected, the *uns-1* mutant showed an increased steady state RNA levels for both *CBF1* and *SVK* in response to cold whereas the *cbf1-3* mutant displayed a substantially decreased induction of *CBF1* and increased *SVK* levels (Fig. 1C-D). The *svk-1* mutant showed a decreased *SVK* level and a stronger induction of *CBF1* compared to wild-type. This data supports our initial hypothesis that limited transcription on one strand can decrease the frequency of the collision events and increase the number of transcription events on the other strand to reach their preferred polyadenylation site. Thus, the restriction of transcription in a strand specific manner in the T-DNA lines not only reinforces collision mechanism at the *CBF1-SVK* regulatory circuit, but also corroborate with the collision events detected by plaNET-seq.

**Figure 1:**
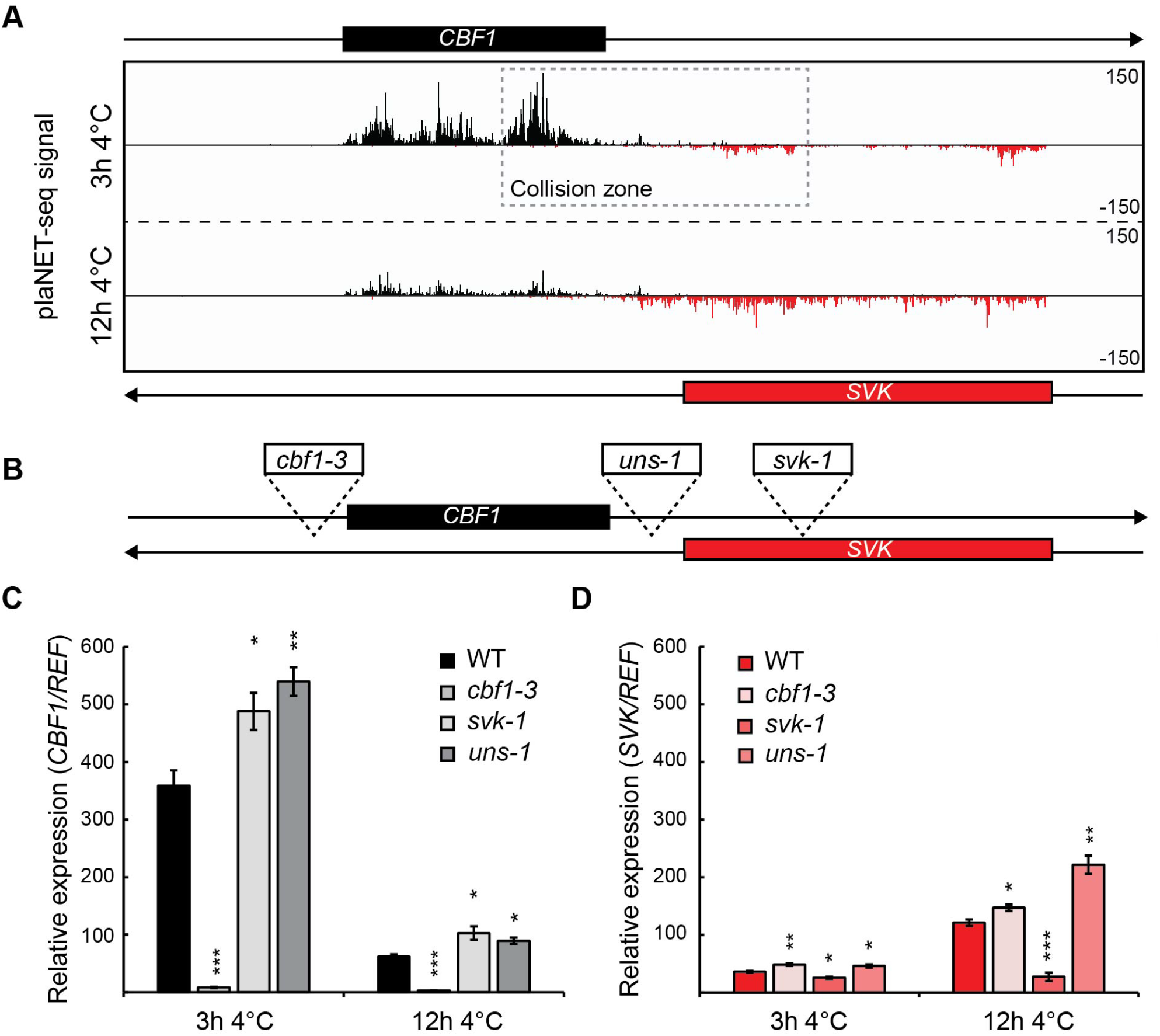
RNAPII collisions can be inferred by restricting expression strand-specifically. A planet-seq signal after 3 hours and 12 hours at 4°C at the *CBF1* and *SVK* locus. Nascent RNAPII transcription is shown for sense and antisense transcripts in black and red, respectively. B Graphical representation of the insertion positions of the T-DNA lines used in this study. C, D Relative *CBF1* (C) and *SVK* (D) expression determined by RT-qPCR in WT and mutants that affect the expression of *CBF1* and *SVK* during cold exposure. Bars represent mean ± SEM from three biological replicates. The relative level of *CBF1* and *SVK* transcripts were normalized to the level in WT in control conditions. Statistically significant differences between means were calculated with Student’s t-test (*p < 0.05, **p < 0.01, ***p < 0.001).

### A long non-coding RNA, *SVALNA*, is transcribed downstream of *CBF3*

Interestingly, we detected a similar situation to *SVK-CBF1* at the neighboring *CBF3* locus. Here, a long non-coding RNA (hereafter named *SVALNA, SVN*) was detected by plaNET-seq (Fig. 2A). Similarly to *SVK, SVN* is upregulated by cold and reaches higher transcriptional activity later in the cold response compared to *CBF1* and *CBF3*. We also detected RNAPII stalling along both strands after 3h of 4°C at the *SVN-CBF3* locus (Fig. 2A). However, in contrast to *SVK, SVN* is barely detected in wild-type but clearly accumulates in a nuclear exosome mutant, *hua enhancer 2-2* (*hen2-2*) (Fig. 2B). This suggests that *SVN* is a target for rapid degradation by the nuclear exosome and further indicates that *SVN* transcription is more important than the *SVN* RNA product. To characterize transcription events along *SVN*, we used available transcription start sequencing data in wild-type and *hen2-2* (Thomas *et al*, 2020) to detect the main initiation site (1124 bp downstream of the poly(A) site of *CBF3*) (Supplemental Fig. 1A). With available direct RNA-sequencing data (Schurch *et al*, 2014), we detected two clusters of end sites for *SVN* (Supplemental Fig. 1A), which corresponded to the two isoforms (514 bp and 1103 bp) detected by our Northern blot analysis (Fig. 2B, Supplemental Fig. 1A). The longer isoform was the dominant one at 8h at 4°C compared to earlier in the cold response (Fig. 2B) and its poly(A) site was only 21 bp from the main poly(A) site of *CBF3* (Supplemental Fig. 1A). We confirmed the expression pattern of *SVN* and its nuclear exosome sensitivity with RT-qPCR (Fig. 2C). The *sop2-1* allele has a point mutation in the RRP4 exosome subunit and similarly to *hen2-2*, the mutant accumulated *SVN*. Further confirmation could be obtained by publicly available RNA-seq data from wild-type and *hen2-2* (Supplemental Fig. 1B). Taken together, *SVN* is a long non-coding RNA transcribed downstream of *CBF3* on the antisense strand. In contrast to the well-described *SVK, SVN* is a nuclear exosome target and rapidly degrades in wild-type plants. Further, *SVN* has a potential regulatory role of *CBF3* based on RNAPII stalling patterns along the *SVN-CBF3* region.

**Figure 2:**
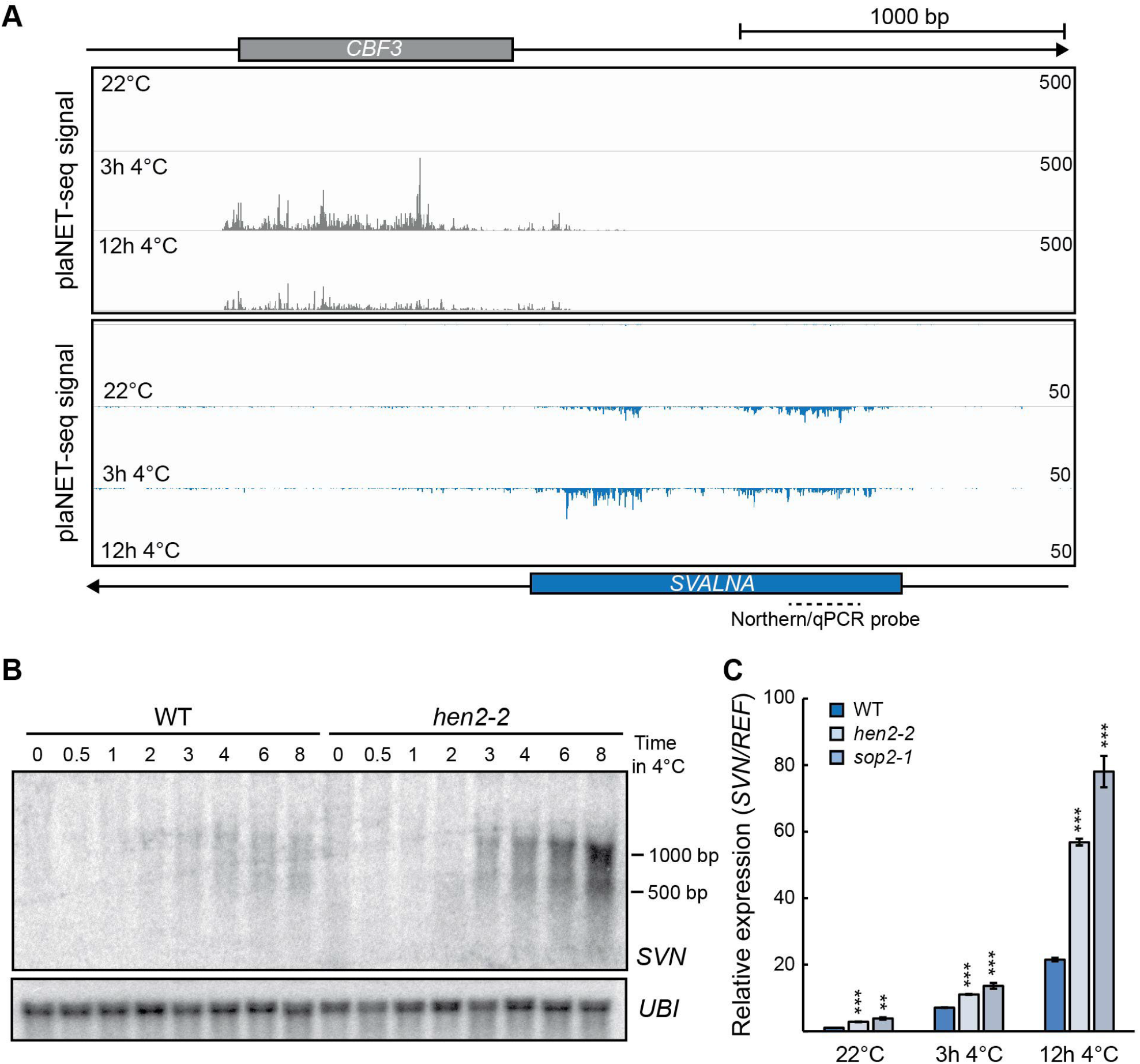
Characterization of the lncRNA *SVALNA*. A planet-seq signal after 3 hours and 12 hours at 4°C and in control conditions at the *CBF3* and *SVN* locus. Nascent RNAPII transcription is shown for sense and antisense transcripts in grey and blue, respectively. The binding site of the northern/qPCR probe is shown. B Northern blot of a cold exposure time series of Col-0 (WT) and *hen2-2*. The probe used for *SVN* is shown in A). Blots were repeated with three biological replicates. *UBI* was used as loading control. C Relative *SVN* expression determined by RT-qPCR in WT and mutants that affect the expression of *SVN* during cold conditions. Bars represent mean ± SEM from three biological replicates. The relative level of *SVN* transcripts were normalized to the level in WT in control conditions. Statistically significant differences between means were calculated with Student’s t-test (*p < 0.05, **p < 0.01, ***p < 0.001).

### *SVN* negatively regulates *CBF3*

To investigate if *SVN* could regulate *CBF3* in a manner similar to the collision mechanism of the *SVK-CBF1* circuit, we isolated a T-DNA line in the *CBF3* promoter (*cbf3-2*) and one line in the 5′-UTR (*cbf3-1*) (Fig. 3A). Unfortunately, no T-DNA lines were available at the *SVN* locus. Therefore, we employed a CRISPR-Cas9 deletion strategy to remove the TSS and part of the promoter of *SVN*. Two independent lines were generated, both deleting the *SVN* TSS and part of the promoter (Fig. 3A, Supplemental Fig. 2A). As expected, the *cbf3* mutants both showed a decreased *CBF3* level after exposure to cold (Fig. 3B). This correlated to an increase of *SVN* levels (Fig. 3C), indicating that, indeed, RNAPII collisions are responsible for the regulation of *CBF3* and *SVN* levels in cold conditions. Deletion of the *SVN* promoter in the *svn* mutants resulted in increased *SVN* and decreased *CBF3* steady state levels (Fig. 3B), a corroborating result of the RNAPII collision mechanism. Interestingly, the activation of *SVN* is correlated to changes in the chromatin environment, specifically around the promoter which is shown in a publicly available ChIP-seq data set (Supplemental Fig. 2B). In this data set a peak of the repressive mark, H3K27me3, could be detected at control conditions. This peak is rapidly decreased in amplitude following cold treatment while the activation mark, H3K4me3 increases (Supplemental Fig. 2B). In summary, *SVN* negatively regulates *CBF3* in a similar manner to *SVK* and *CBF1*. Similarly to *SVK, SVN* switch poly(A) site during prolonged cold exposure to allow transcribing RNAPII complexes in close proximity to the 3′-end of *CBF3*. In contrast to *SVK, SVN* transcription only initiates from a single site, the *SVN* transcript is not spliced and is rapidly degraded by the nuclear exosome throughout the cold exposure. Furthermore, the activation of *SVN* is dependent on chromatin remodeling around its TSS.

**Figure 3:**
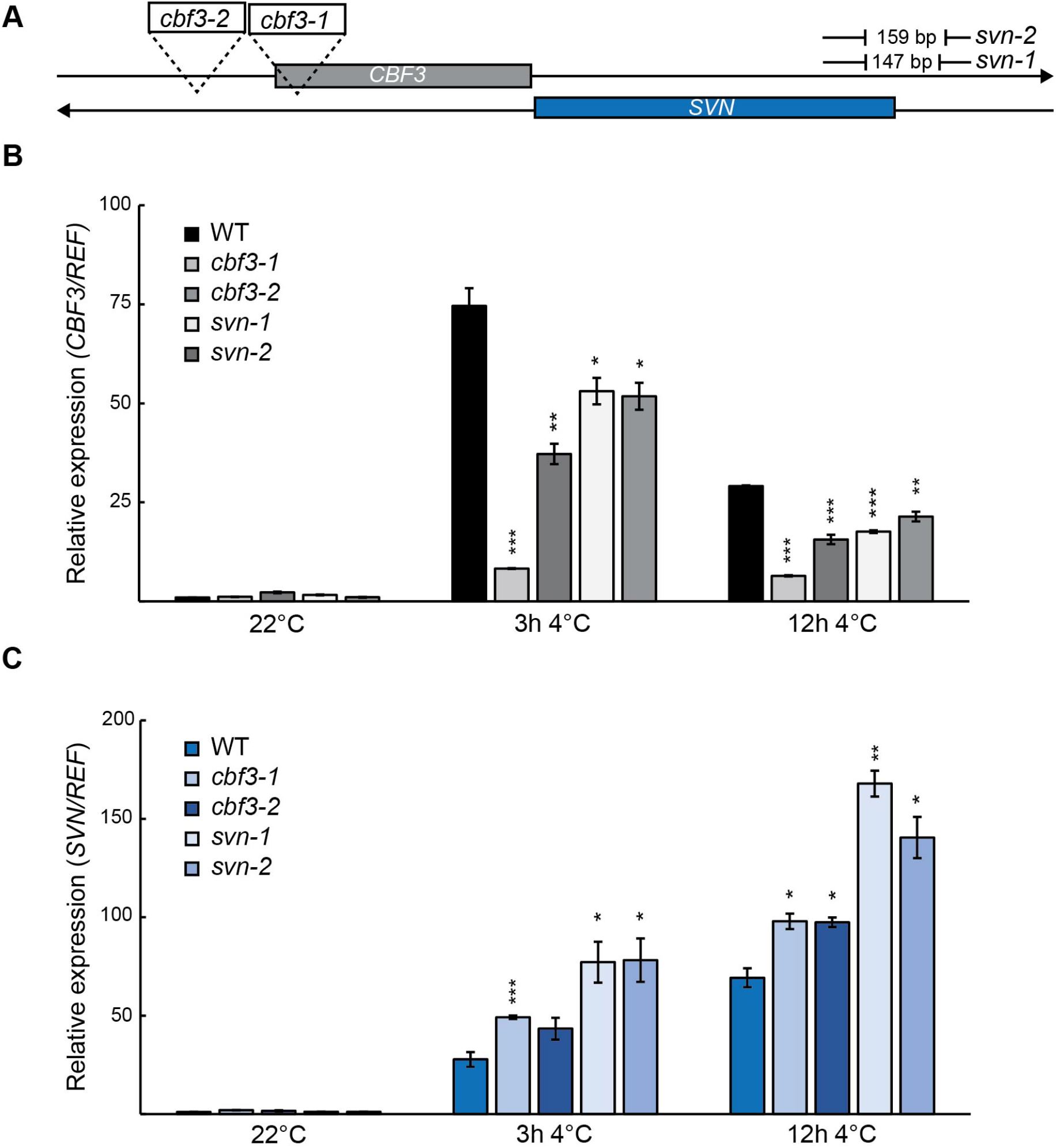
RNAPII collisions occur at the *CBF3-SVN* locus. A Graphical representation of the insertion positions of the T-DNA lines and deletions induced by CRISPR-Cas9 used in this study. B, C Relative *CBF3* (B) and *SVN* (C) expression determined by RT-qPCR in WT and mutants that affect the expression of *CBF3* and *SVN* during cold conditions. Bars represent mean ± SEM from three biological replicates. The relative level of *CBF3* and *SVN* transcripts were normalized to the level in WT in control conditions. Statistically significant differences between means were calculated with Student’s t-test (*p < 0.05, **p < 0.01, ***p < 0.001).

### *SVK* switches transcription start site in prolonged cold exposure

The rapid degradation of *SVN* raises the question of how *SVK* achieves relatively higher stability. We have earlier identified two TSS and two poly(A) signals for *SVK* (Kindgren *et al*., 2018; Zacharaki *et al*., 2023). A group of shorter transcripts (*SVK* α) is initiated from a proximal TSS (in relation to *CBF1*) (Fig. 4A) while transcripts from a distal TSS (*SVK* β) are either terminated at a distal poly(A) site or undergo splicing and continue to the proximal poly(A) site (Fig. 4A). Both *SVK* α transcripts and spliced *SVK* β have a length of around 750 bp, making them difficult to separate on a northern blot with a probe annealing to the 3′-end of *SVK* (Kindgren *et al*., 2018). Thus, to disentangle the active forms of *SVK*, we designed specific RT-qPCR oligos that amplify only *SVK* α or *SVK* β (Fig. 4A). In addition, we designed oligos that would amplify all *SVK* transcripts (Total). In available RNA-seq data, it was clear that the proximal TSS was more active early in the cold response (3h 4°C) while the distal TSS was activated later (12h 4°C) (Fig. 4B), corroborating earlier studies (Kindgren *et al*., 2018). These results were further confirmed by our RT-qPCR analysis (Fig. 4D-F). To investigate if *SVK* transcripts were rapidly degraded by the nuclear exosome, such as *SVN*, we used *sop2-1* and *hen2-2* exosome mutants. We observed that only the *SVK* α transcripts were accumulating in the mutants (Fig. 5A-B), leading to an overall accumulation of *SVK* transcripts (Fig. 5C). The accumulation of *SVK* transcripts only occurred after 12h 4°C, indicating that most amplicons represent unspliced forms of *SVK* β. Combined, these results indicate that *SVK* switches TSS, from a proximal to a distal start site during the cold response. Additionally, the introduction of a splicing event of transcription from the distal TSS may be important for *SVK* function and stability.

**Figure 4:**
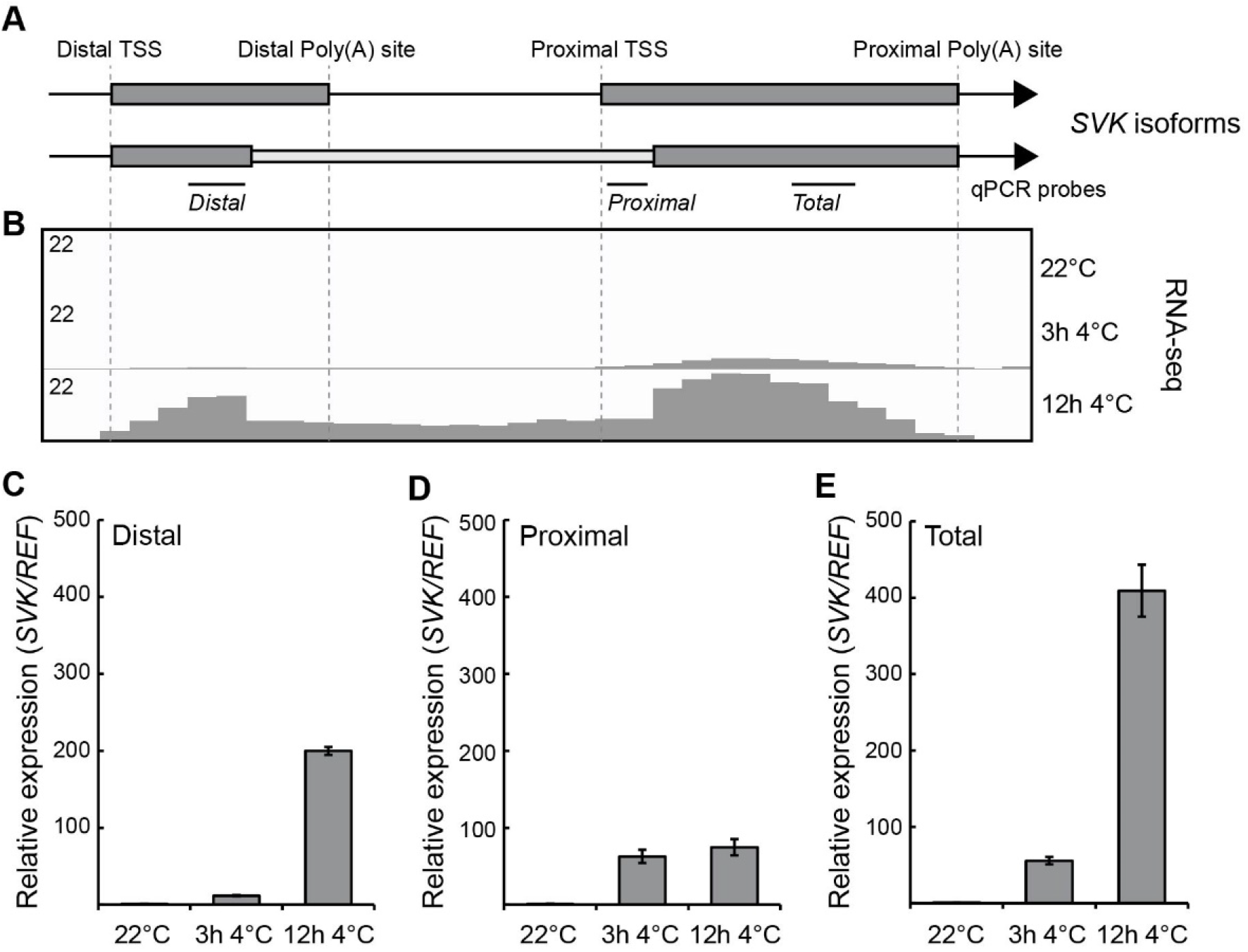
Characterization of *SVK* isoforms. A Graphical representation of *SVK* isoforms and positions of qPCR probes used to identify relative expression of isoforms. B Screenshot of the *SVK* locus from an RNA-seq data set. Elevated transcriptional activity is indicated by higher peak density and amplitude. C-E Relative expression of *SVK* isoforms determined by RT-qPCR in WT during cold exposure. The distal (C), proximal (D) and total (E) transcript of *SVK* were analyzed. Bars represent mean ± SEM from three biological replicates. The relative level of *SVK* transcripts were normalized to the level in control conditions. Statistically significant differences between means were calculated with Student’s t-test (*p < 0.05, **p < 0.01, ***p < 0.001).

**Figure 5:**
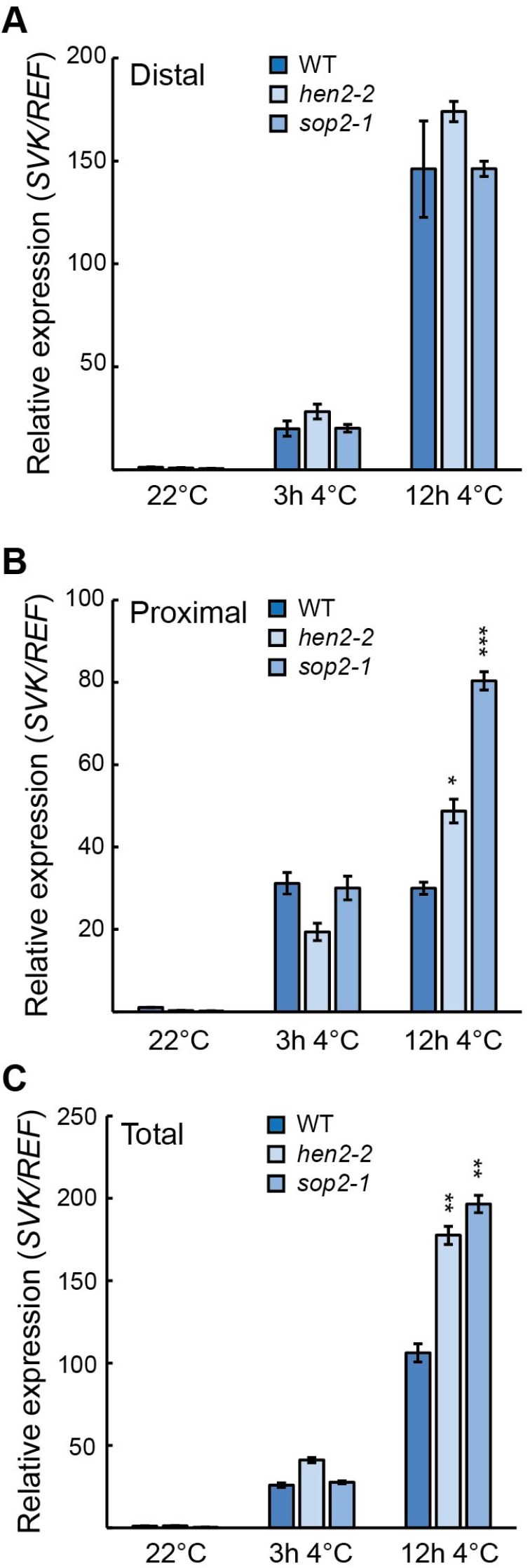
Degradation of *SVK* isoforms. A-C Relative expression of *SVK* isoforms determined by RT-qPCR in WT and exosome mutants that affect the expression of *SVK* during cold conditions. The distal (A), proximal (B) and total (C) transcript of *SVK* were analyzed. Bars represent mean ± SEM from three biological replicates. The relative level of *SVK* transcripts were normalized to the level in WT in control conditions. Statistically significant differences between means were calculated with Student’s t-test (*p < 0.05, **p < 0.01, ***p < 0.001).

### Splicing of *SVK* is essential for its *trans*-acting role

To test the involvement of splicing in the maturation of *SVK*, we used two mutants of the spliceosome, *sm-like 7-1* and *8-1* (*lsm7-1* and *lsm8-1*). Both mutants are part of the nucleoplasmic version of the spliceosome (Nardeli *et al*, 2023; Perea-Resa *et al*, 2012). We used longer exposure to cold in these experiments (12 and 24h 4°C) to investigate the reported *trans*-role for *SVK* on *CBF3* (Gómez-Martínez *et al*., 2023). In addition, we used qPCR oligos to encompass spliced and unspliced versions of *SVK* (Fig. 6A). The spliced version of *SVK* was greatly reduced in the two mutants while the unspliced version was increased (Fig. 6B-C). This splicing mis-regulation of *SVK* had a significant effect on the repressive role of *CBF3* (Fig. 6D), indicating that splicing is essential for the *trans* role of *SVK*. These results suggest that the splicing machinery is crucial for the mode-of-action of *SVK* on *CBF3*.

**Figure 6:**
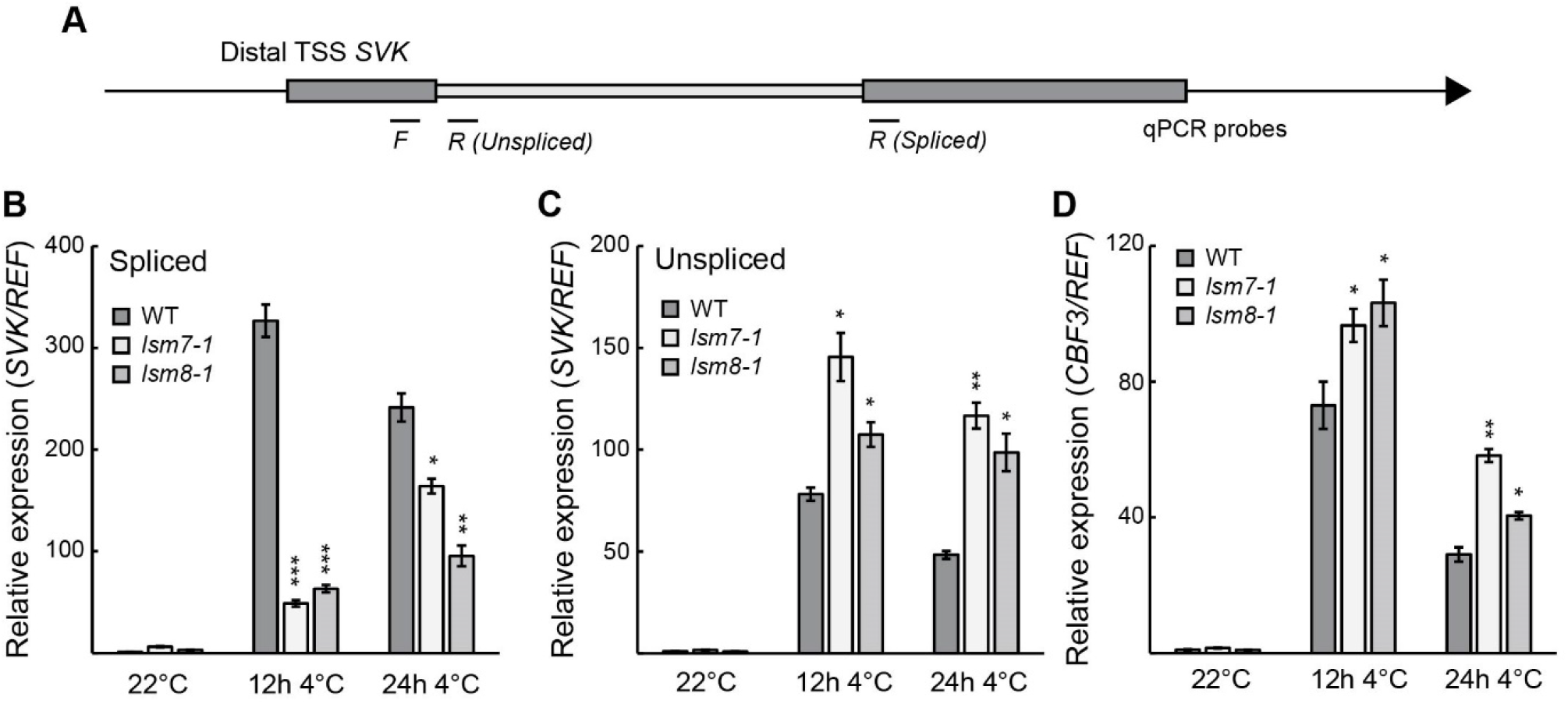
Splicing of *SVK*. A Graphical representation of *SVK* β and the positions of qPCR probes used to identify relative expression of spliced and unspliced isoforms. B-D Relative expression of the spliced *SVK* (B) and unspliced *SVK* (C) isoforms and *CBF3* (D) determined by RT-qPCR in WT and mutants that affect the expression of *SVK* and *CBF3* during cold exposure. Bars represent mean ± SEM from three biological replicates. The relative level of *SVK* transcripts were normalized to the level in WT in control conditions. Statistically significant differences between means were calculated with Student’s t-test (*p < 0.05, **p < 0.01, ***p < 0.001).

### Activation of the distal TSS of *SVK* is important for proper *CBF1* and *CBF3* regulation

Our results so far point to an important switch from a proximal to distal TSS for *SVK* and an introduction of a splicing event of *SVK* transcripts initiating from the distal TSS to surveille the levels of *CBF1* and *CBF3*. To further show the significance, we obtained CRISPR-Cas9-mediated deletion lines targeting the promoter of the distal TSS. A stable line designated as *svk-2*, with a 32 bp deletion, 66 bp upstream of the distal TSS was generated (Fig. 7A, Supp. Fig. 3A). In *svk-2*, only the distal transcription was affected (Fig. 7B-C). This leads to an overall lower *SVK* level after 12h 4°C (Fig. 7D). Furthermore, the peak of *CBF1* and *CBF3* expression (3h at 4°C) was not mis-regulated in *svk-2*, but the repression of both genes (12h at 4°C) was (Fig. 7E-F). Thus, the transcription from the distal TSS of *SVK* is crucial for its repressive role on *CBF1* and *CBF3*. Interestingly, the promoter of the distal TSS is heavily decorated with repressive H3K27me3 marks at control conditions (Supp. Fig. 3B), similar to what was observed around the *SVN* promoter. The H3K27me3 signal decreases after cold treatment, opening up the regulatory sequences of the distal TSS and transcription is initiated, as seen by an increased H3K4me3 signal (Supp. Fig. 3B). In summary, *SVK* function requires several steps of transcriptional regulation such as TSS switch and splicing, as well as remodeling of the chromatin environment. All in all, our study highlights the mechanistically complex and active regulation of two long non-coding transcripts, *SVK* and *SVN*, to fulfill their regulatory role on coding *CBF1* and *CBF3* mRNA levels for proper cold adaptation in Arabidopsis.

**Figure 7:**
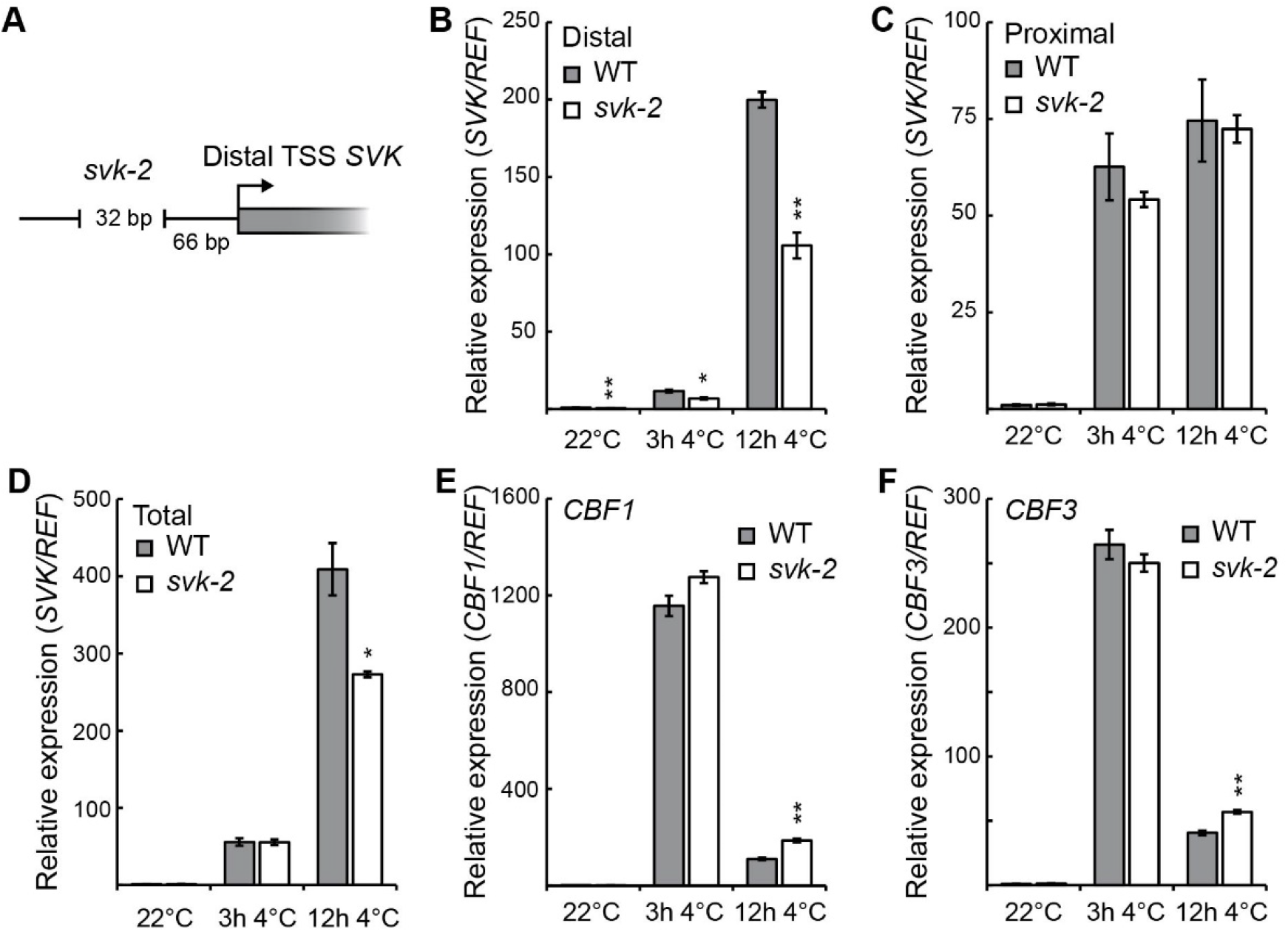
Characterization of *SVK* CRISPR-Cas9 deletion line *svk-2*. A Graphical representation of the position of the deletion in the line *svk-2* induced by CRISPR-Cas9 used in this study. B-D Relative expression of *SVK* isoforms determined by RT-qPCR in WT and *svk-2* that affect the expression of *SVK* during cold conditions. The distal (B), proximal (C) and total (D) transcript of *SVK* were analyzed. Bars represent mean ± SEM from three biological replicates. The relative level of *SVK* transcripts were normalized to the level in WT in control conditions. Statistically significant differences between means were calculated with Student’s t-test (*p < 0.05, **p < 0.01, ***p < 0.001). E, F Relative *CBF1* (E) and *CBF3* (F) expression determined by RT-qPCR in WT and *svk-2* that affect the expression of *CBF1* and *CBF3* during cold conditions. Bars represent mean ± SEM from three biological replicates. The relative level of *CBF1* and *CBF3* transcripts were normalized to the level in WT in control conditions. Statistically significant differences between means were calculated with Student’s t-test (*p < 0.05, **p < 0.01, ***p < 0.001).

## DISCUSSION

Here, we present an updated model (Fig. 8) showing how the long non-coding RNAs *SVK* and *SVN* regulate *CBF1* and *CBF3*. Under normal conditions (22°C), *CBF* and *SVK* transcription is low, with the long *SVK* α isoform forming double-stranded RNA with *CBF1* mRNA, leading to cleavage by AGO1. In contrast, *SVN* is rapidly degraded by the nuclear exosome. During initial cold exposure (3 - 12 hours), transcription of *CBF* genes and the lncRNAs *SVK* and *SVN* increases, causing premature mRNA termination of *CBF1* and *CBF3* due to RNAPII collisions. A switch in TSS results in *SVK* β becoming dominant later in the cold response. Maturation in the form of *SVK* β splicing produces a stable transcript that recruits the PRC2 complex, adding repressive marks on *CBF3* to suppress its expression. In summary, this model highlights the importance of lncRNA isoforms and expression patterns in regulating *CBF1* and *CBF3*.

**Figure 8:**
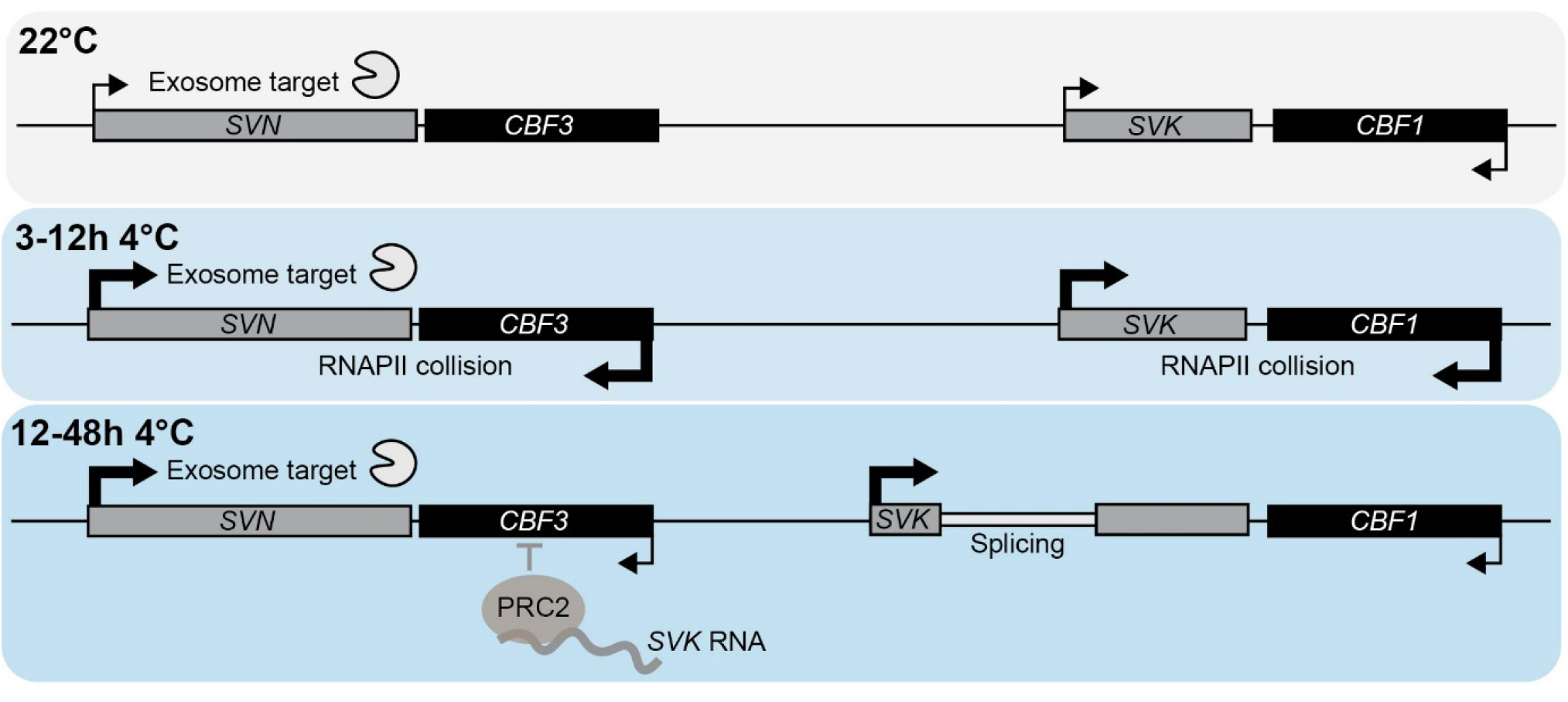
Model of how lncRNAs *SVK* and *SVN* regulate expression of *CBF1* and *CBF3*. At 22°C, *CBF* and *SVK* transcription levels are low, with the long *SVK* α isoform forming double-stranded RNA with *CBF1* mRNA, following degradation. *SVN* is quickly degraded by the nuclear exosome. Repressive H3K27me3 marks are present at both loci. Exposure to cold for 3 to 12 hours boosts transcription of *CBF* genes and *SVK* and *SVN*, causing premature termination of *CBF1* and *CBF3* mRNA due to RNA polymerase II collisions. Immature mRNA is degraded. During later cold response, *SVK* β becomes the dominant isoform. Splicing of *SVK* β produces a stable transcript that recruits the PRC2 complex, adding repressive marks to *CBF3* and reducing its expression.

A central theme in our findings is the role of histone modifications, particularly the dynamic levels of repressive H3K27me3 in the *CBF* region. H3K27me3 marks are evicted during cold exposure around the TSS of *SVN* and the distal TSS of *SVK*. This decrease in repressive marks is a well-documented epigenetic mechanism that allows plants to activate genes in response to environmental stimuli (Shen *et al*, 2021; Yuan *et al*, 2013). Interestingly, only the distal TSS of *SVK* is decorated with high levels of H3K27me3 at control conditions, not the proximal TSS, explaining the TSS switch to produce *SVK* β when H3K27me3 is removed during the later stages of the cold response. This, in turn, results in PRC2-mediated H3K27me3 enrichment over the *CBF3* gene body by *SVK* β (Gómez-Martínez *et al*., 2023). While the players involved in removing H3K27me3 from the TSS of *SVK* and *SVN* are unknown, there is an established link between lncRNA and PRC2 to add repressive marks to histones. PRC2 regulates numerous targets throughout the genome yet lacking intrinsic DNA-binding affinity. To achieve specificity, PRC2 can form subcomplexes with polycomb-like proteins (PCLs) which help recruit it to unmethylated CpG islands recognized by PCLs (Li *et al*, 2017). Additionally, DNA motifs such as GA-repeats, telobox-like motifs and RY motifs have been shown to assist PRC2 to specific genes (Qüesta *et al*, 2016; Shu *et al*, 2020; Wu *et al*, 2020; Xiao *et al*, 2017; Yuan *et al*, 2016; Zhou *et al*, 2018). Moreover, the interaction between lncRNAs and PRC2 is emerging as a crucial element for balancing gene silencing and activation in eukaryotes (Brockdorff, 2013; Zhao *et al*., 2018). The role of the PRC2-lncRNA complexes extend beyond a singular case in Arabidopsis; the lncRNAs *COLDAIR* and *COLDWRAP* bound to PRC2 are essential for the repression of the *FLOWERING LOCUS C* (*FLC*) gene during the vernalization process. (Heo & Sung, 2011; Kim & Sung, 2017; Kim *et al*., 2017). Importantly, the PRC2-lncRNA regulatory complex is conserved in other eukaryotes. For example, the human lncRNA *HBL1* interacts with PRC2 in early cardiogenesis by guiding PRC2 occupancy on crucial cardiogenic genes in pluripotent stem cells (Liu *et al*, 2021). Similarly, the lncRNA *Gm15055* in mouse embryonic stem cells represses the expression of the Hoxa gene by recruiting PRC2 to maintain H3K27me3 at the locus (Liu *et al*, 2016). In the case of the lncRNA *LEVER* in human cells, the non-polyadenylated, nascent RNA produced by the locus binds and inhibits PRC2 function of the adjacent *β-globin* gene (Teo *et al*, 2022). The promiscuous binding of RNA to PRC2 might be an important regulatory point for the addition/removal of H3K27me3 at specific gene loci depending on what RNAs are produced in the vicinity and an avenue for future research. Findings from various eukaryotic organisms underscore a general role for lncRNAs in modulating PRC2 activity, highlighting their importance in the broader context of epigenetic regulation and gene expression control in eukaryotes.

The regulation of the PRC2-*SVK* complex on *CBF3* requires splicing of *SVK*. Thus, the bound transcript (*SVK* β) must possess specific features enabling that interaction. Modular stem-and-loop structures within the RNA have been shown to form unique motifs that allow lncRNAs to bind to proteins (Guttman & Rinn, 2012; Kim *et al*., 2017; Mercer & Mattick, 2013; Somarowthu *et al*, 2015). For instance, *HOTAIR*, a well-studied PRC2 interacting lncRNA in humans, folds into a complex secondary structure that facilitates binding to proteins (Somarowthu *et al*., 2015). Amongst others, the lncRNA *COOLAIR* which forms a complex structure with multi-helix junctions was shown to have an evolutionary conserved secondary structure, making lncRNA transcripts important regulators of gene expression (Corona-Gomez *et al*, 2023; Hawkes *et al*, 2016). RNA processing such as splicing is critical for the functional role of lncRNAs. The lncRNA *HOTAIR* produces three different splice variants. The 5′ end of its longest isoform acts as a scaffold to bind chromatin-modifying complexes (Khan *et al*, 2023; Tsai *et al*, 2010; Wu *et al*, 2013). However, another isoform, generated by alternative splicing, lacks the binding scaffold (Loewen *et al*, 2014), enabling *HOTAIR* to differentially modulate target genes (Khan *et al*., 2023). This process emphasizes the pivotal role of alternative splicing of lncRNAs in plants. Kiegle et al. (2018) characterized the lncRNA *LOC9270896* and its isoforms, demonstrating that intron retention and exon skipping yield multiple transcripts crucial for rice seed maturation (Kiegle *et al*, 2018). The splicing process generates diverse isoforms that can affect RNA stability, localization, and interaction with other molecular partners, thus modulating gene expression (Deveson *et al*, 2018; Kalsotra & Cooper, 2011; Kiegle *et al*., 2018). Our findings indicate that the splicing of *SVK* β is crucial for down-regulating *CBF3* following extended cold exposure as evidenced by measuring gene expression levels in spliceosome mutants (Fig. 6). A decrease in spliced transcript levels leads to elevated *CBF3* expression during cold stress, affirming the significance of the spliced transcript in gene regulation. This raises questions regarding the evolutionary pressures that shape the functional diversity of lncRNAs. Unlike antisense lncRNAs that interact via base-pairing (Meena *et al*, 2024), non-antisense lncRNAs like *SVK* can serve as scaffolds or guides for chromatin modifiers (Gómez-Martínez *et al*., 2023). They can regulate gene expression in *cis* through transcriptional interference, leading to collisions between RNAPIIs (Kindgren *et al*., 2018). The stability of *trans*-acting lncRNA isoforms, particularly *SVK*’s stable forms resulting from the use of alternative transcription start sites and splicing, is crucial for its regulatory function. This aligns with findings that stable lncRNAs effectively regulate gene expression, as seen with the mammalian lncRNA *FIRRE*, which has both *cis* and *trans* effects (Fang *et al*, 2020) and in Arabidopsis where *APOLO* regulates the gene *PID* in *cis* and the *PID* homolog *WAG2* in *trans* by forming R-loops (Ariel *et al*., 2014; Ariel *et al*, 2020). The post-transcriptional regulation of lncRNAs adds another layer of complexity for how their function can be controlled to interact with the correct partners at the right time during stress response or development.

In conclusion, our study elucidates the pivotal role of lncRNAs *SVK* and *SVN* in the regulation of *CBF1* and *CBF3* gene expression in response to cold stress. We present an updated model demonstrating that through several mechanisms such as changes in transcription dynamics by RNAPII stalling caused by a switch in isoforms, alternative splicing and recruiting PRC2 complex as well as histone modifications, lncRNAs are crucial in regulating gene expression. The structural characteristics of *SVK* β are essential for its interactions and regulatory functions, emphasizing how splicing generates diverse isoforms that fine-tune gene expression during environmental challenges. Our findings not only advance our understanding of lncRNA functionality but also highlight their fundamental importance as key players in networks that govern gene expression across diverse biological contexts. Ultimately, lncRNAs are not merely regulatory elements, they are the cornerstone of complex gene regulation, shaping the biological responses of organisms to their environments.

## METHODS AND MATERIAL

### Plant growth and conditions

For the wild-type background Arabidopsis (*Arabidopsis thaliana*) Col-0 or Columbia accession was employed. For the growth of plants, seeds were surface sterilized and stratified for 2-4 days at 4° C in the dark and either transferred to soil directly or plated on ½ Murashige and Skoog (MS) basal medium supplemented with 1% (w/v) sucrose. Plants were grown in long day conditions (16h light, 8 dark, ∼100 µE), SciWhite LEDs (Percival) for 10 days. Biological replicates in all experiments represent approximately 20-30 seedlings grown on separate plates. Cold treatment (4° C, ∼25 µE) was initiated at ZT4. T-DNA insertional mutant lines used have been described elsewhere: *svk-1* (GABI_145A05) and *uns-1* (SALK_018442) (Kindgren *et al*., 2018), *cbf3-1* (SAIL_244_D02) (Khanna *et al*, 2006), *lsm7-1* (SALK_066076) (Nardeli *et al*., 2023), *lsm8-1* (SALK_048010) (Perea-Resa *et al*., 2012). The *cbf3-2* (GABI_136G01) mutant was isolated in this study and ordered from the Nottingham Arabidopsis Stock Centre. All T-DNA lines were genotyped and confirmed for homozygosity by PCR. Oligos for genotyping can be found in Supp. Table 1.

### RNA extraction, cDNA synthesis and RT-qPCR

Total RNA extraction from plant material was carried out using Plant RNA Mini Kit (Omega-Biotek) as per suppliers’ instructions. 1 µg of the extracted RNA was additionally treated with dsDNase (Thermo Fisher Scientific) for the elimination of genomic DNA contamination. Successively, complementary DNA (cDNA) synthesis was carried out using iScript (BioRad) reverse transcriptase as per manufacturer instructions together with a reference gene. Quantitative real-time PCR (RT-qPCR) was performed on CFX384 Real-Time PCR detection systems (BioRad) using SYBR premix (BioRad), cDNA, reverse primer and forward primers at the concentration of 10 pmol/µl with the PCR cycler following initial denaturation at 95 ° C for 30s, standard 40 cycles of 94 ° C for 10 s, 60 ° C for 30s. The specificity of RT-qPCR products was assessed from the single peak melt curves. For the data analysis, the Cq values from a minimum of 3 biological replicates with 3 technical replicates were averaged and △Cq was obtained as Cq (gene of interest)-Cq (reference gene). Final calculations were performed by following with 2^(-Δcq) or 2^(-ΔΔcq), adjusted to experimentally determined primer efficiency for determination of fold change in gene expression levels. Statistically significant differences were calculated with Student’s t-test. Primers used are listed in Supp. Table 1.

### Northern blotting

Total RNA was extracted using RNeasy Plant Mini Kit (Qiagen, Germany) according to manufacturer’s instructions. 20 µg of total RNA was separated on a 1.2% agarose gel with formaldehyde and 1xMOPS. Gels were blotted overnight onto a nylon membrane and crosslinked with UV radiation. Probes were made in a PCR reaction by incorporating radioactive dTTP (32P, PerkinElmer, USA) from a DNA probe template. Membranes were exposed to a phosphorimager screen (GE Healthcare, UK) for 1-10 days depending on the expression level of the transcript of interest. Screen was subsequently scanned with a Typhoon scanner (GE Healthcare, UK).

### Generation of CRISPR-Cas9 deletion lines

Guide RNAs (gRNAs) for the generation of CRISPR-Cas9 deletion lines *svk-2, svn-1* and *svn-2* were designed using the CHOPCHOP webserver (http://chopchop.cbu.uib.no/). A 2gRNA fragment was amplified utilizing the DT1T2 plasmid (Xing *et al*, 2014) using Phusion DNA polymerase (Thermo Fisher Scientific). The oligonucleotides used are listed in Supp. Table 1. The resulting PCR product was separated on a gel, excised, purified, and cloned into the targeting vector CF588 (kindly provided by Christian Fankhauser). The modified CF588 vector contains a GFP seed coat expression cassette to enable faster screening. Final plasmids were validated by sequencing and introduced into WT Col-0 plants via *Agrobacterium tumefaciens* (GV3101) floral dip. T1 seeds were initially screened for GFP expression and subsequently genotyped by PCR. In the T2 generation, seeds lacking GFP fluorescence were selected, and homozygous plants were confirmed by PCR and sequencing for genomic deletion. Seeds from confirmed homozygous plants were used for subsequent experiments.

### Genome-wide data sets

Available genome-wide datasets used in this study include plaNETseq (Kindgren *et al*., 2020), RNA-seq (Bhat *et al*, 2024), TSS-seq (Thomas *et al*., 2020), DRS-seq (Schurch *et al*., 2014), and H3K27me3 and H3K4me3 ChIP-seq (Faivre *et al*, 2024).

## ACKNOWLEDGEMENTS

We would like to thank members of the Kindgren lab for their critical reading of the manuscript. We would like to thank the greenhouse personnel at Umeå Plant Science Centre for plant maintenance. We are grateful to Dr. Markus Schmid and Dr. Sarah Nardeli for the gift of *lsm7-1* and *lsm8-1* seeds. We thank Dr. Sebastian Marquardt for initial discussions related to the project. We are grateful for the support from Bio4Energy (https://bio4energy.se). The research in the P.K. lab was funded by the Swedish research council (2018-03926), FORMAS (2021-01065), Kempe Foundation (JCK-3131), and grants from the Knut and Alice Wallenberg Foundation and the Swedish Governmental Agency for Innovation Systems [KAW 2016.0355 and 2020.0240, VINNOVA 2016-00504].

