## Supplemental Figures for "Two long non-coding RNAs, *SVALKA* and *SVALNA*, regulate *CBF1* and *CBF3* via multiple mechanisms"

**SUPPLEMENTAL FIGURES – EXPANDED VIEW**

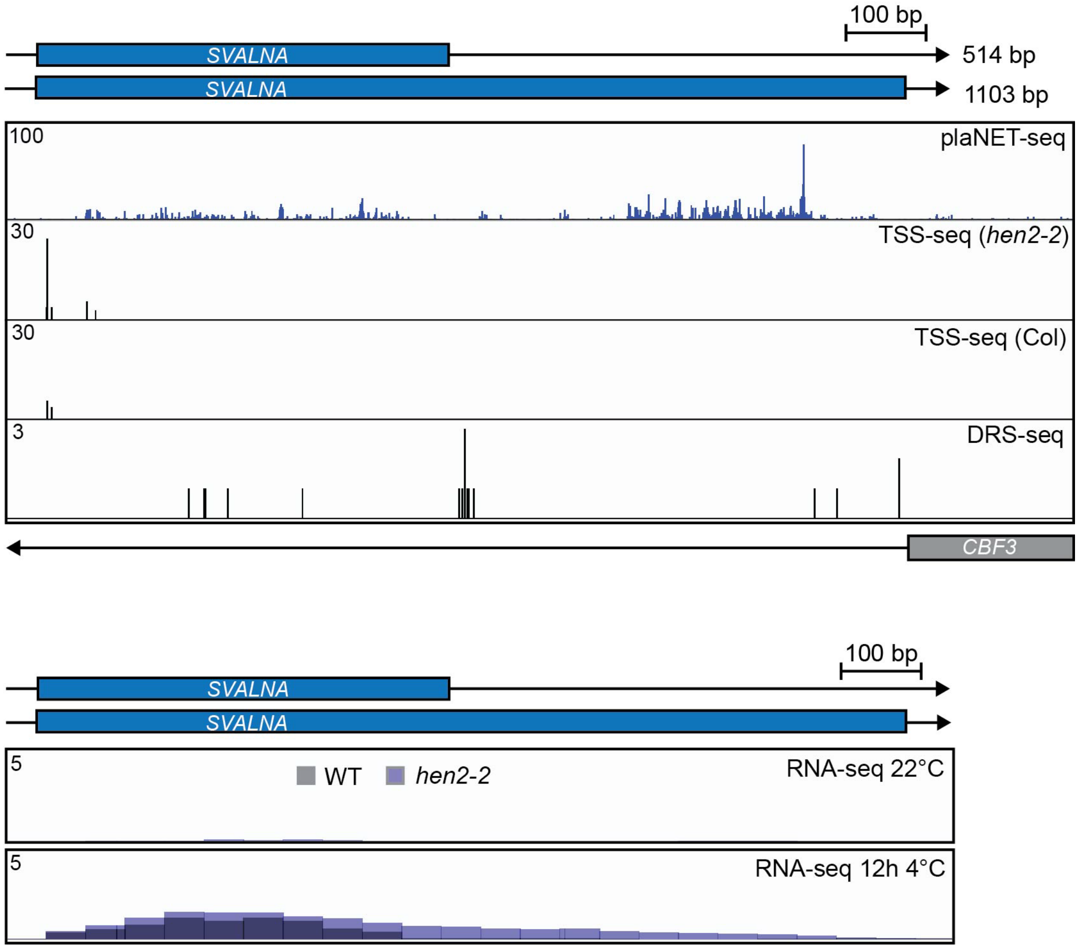


**Supp. Figure 1 - Detection and characterization of *SVN* transcripts.**

A plaNET-seq coverage profile for *SVN* with positions of RNAPII are shown for the sense strand in blue. TSS-seq (transcription start site sequencing) for WT and *hen2-2* and DRS-seq (direct RNA sequencing) for WT tracks are also shown. The DRS-seq track reveals sites of mRNA cleavage and polyadenylation (PAS).

B Screenshot of the *SVN* locus from an RNA-seq data set. Shown are WT and *hen2-2* at 22°C and after 12 hours at 4°C. Elevated transcriptional activity is indicated by higher peak density.

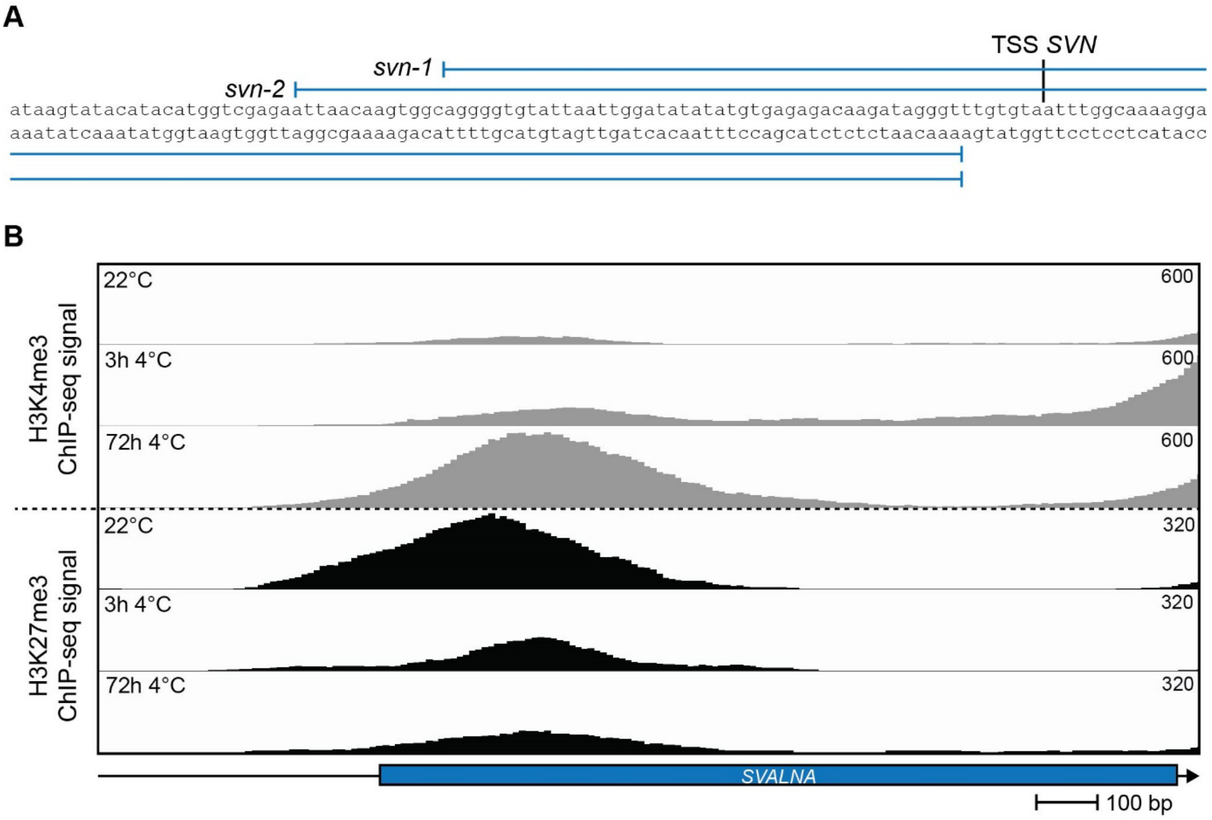


**Supp. Figure 2 – Characterization of *SVN* CRISPR-Cas9 deletion lines *svn-1* and *svn*-2 and histone marks along the gene body of *SVN*.**

A Graphical representation of the position of the deletion in the lines *svn-1* and *svn-2* induced by CRISPR-Cas9 used in this study.

B Screenshot of the *SVN* locus from a ChIP-seq data set. Shown are WT at 22°C and after 3 hours and 72 hours at 4°C. Higher occupancy of H3K4me3 and H3K27me3 are indicated by higher peak density and amplitude.

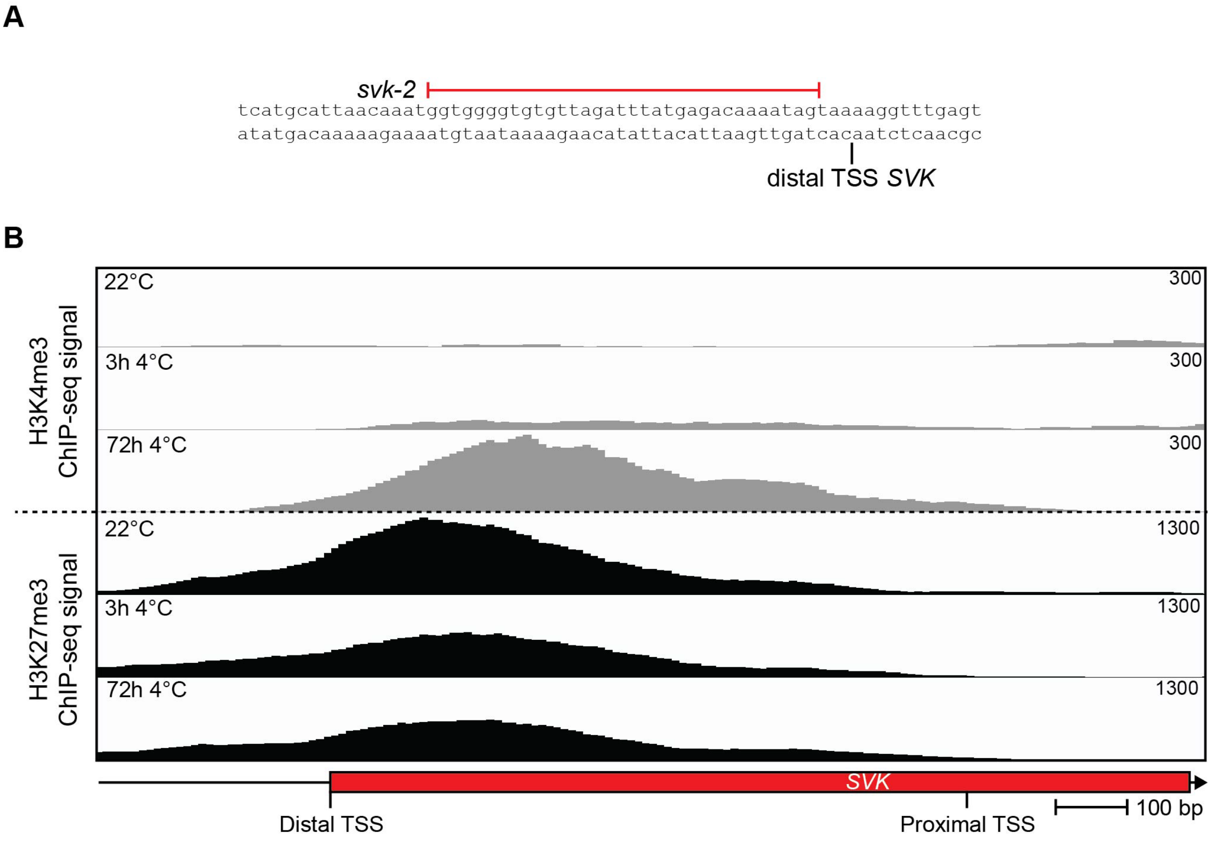


**Supp. Figure 3 - Characterization of *SVK* CRISPR-Cas9 deletion line *svk*-2 and histone marks along the 5′ end of *SVK*.**

A Graphical representation of the position of the deletion in the line *svk-2* induced by CRISPR-Cas9 used in this study.

B Screenshot of the *SVK* locus from a ChIP-seq data set. Shown are WT at 22°C and after 3 hours and 72 hours at 4°C. Higher occupancy of H3K4me3 and H3K27me3 are indicated by higher peak density and amplitude.
