## Supplemental Table for "Two long non-coding RNAs, *SVALKA* and *SVALNA*, regulate *CBF1* and *CBF3* via multiple mechanisms"

**Supplementary Table 1: Oligos used in this study**

### Cloning

| Locus | Other name | Sequence | Comment |
|---|---|---|---|
| At4g07390 | *SVN* | aacaggtctctattGGAGAATTAACAAGTGGCAGgttttagagctagaaatagc<br>aacaggtctctaaacTACTTTTGTTAGAGAGATGCaatctcttagtcgactctac | guide RNAs for the generation of CRISPR-Cas9 deletion line *svn-1* |
| | | aacaggtctctattGATGGTCGAGAATTAACAAGgttttagagctagaaatagc<br>aacaggtctctaaacTACTTTTGTTAGAGAGATGCaatctcttagtcgactctac | guide RNAs for the generation of CRISPR-Cas9 deletion line *svn-2* |
| At4g07395 | *SVK* | aacaggtctctattgaagcatgcAGTTGTAATTGgttttagagctagaaatagc<br>aacaggtctctaaacCACCATTTGTTAATGCATGCaatctcttagtcgactctac | guide RNAs for the generation of CRISPR-Cas9 deletion line *svk-2* |

### Genotyping

| Locus | Mutant | Sequence | Comment |
|---|---|---|---|
| At4g07390 | *svn-1, svn-2* | ACACAAGTTGCTTAAAATCGAAGG<br>AGACAAGTAGCGAAGGGACG | |
| At4g07395 | *svk-2* | GCACGTAAGTCACCAAGTAG<br>AATCATATGACTAAGGACGTGG | |
| At4g25480 | *cbf3-2 (*GABI_136G01*)* | AGTTTCTCCGACGAACTCCTC<br>TTAGCAGTCTATTTAGGGTTTTCC | |
| | | ATATTGACCATCATACTCATTGC | GABI-KAT LB |

### RT-qPCR/Northern

| Locus | Mutant | Sequence | Comment |
|---|---|---|---|
| At4g07390 | *SVN* | TCTCCCCCTTGGCTTAGACT<br>TTTGCCGGAAAACTCAACTC | Amplification of *SVN* levels/Northern probe |
| At4g07395 | *SVK* | acccgaaataaacaaatccgta<br>ggagaagcaagacgacaacg | Amplification of total *SVK* levels |
| | | aaagtctgcctcgttacctac<br>cgttgtcgtcttgcttctcc | Amplification of proximal *SVK* levels |
| | | tttccttcaaccaccacgtc<br>ccttgggaggaaattcttcta | Amplification of distal *SVK* levels |
| | | ttttcttgaccatacccactct<br>tttccttcaaccaccacgtc | Amplification of spliced *SVK* levels |
| | | tgtgatcaaaagggttagcacg<br>cgtctcgaatctagcaacca | Amplification of unspliced *SVK* levels |
| At4g25490 | *CBF1* | tcgatagtcgtttccatttttgt<br>aaaatgaaggaaccattctaaaaa | Amplification of *CBF1* levels |
| At4g25480 | *CBF3* | tttagaatggaatcttcattatgtttg<br>cccacacttatactgaaactgaatc | Amplification of *CBF3* levels |
| At4g05320 | *UBQ10* | GGCCTTGTATAATCCCTGATGAATAAG<br>AAAGAGATAACAGGAACGGAAACATAGT | Reference gene. Amplification of coding *UBQ10* |
| At3g18780 | *ACT2* | CTTGCACCAAGCAGCATGAA<br>CCGATCCAGACACTGTACTTCCTT | Reference gene. Amplification of coding *ACT2* |
| At4g36800 | *UBI* | CTGTTCACGGAACCCAATTC<br>GGAAAAAGGTCTGACCGACA | Reference gene for Northern blot |
